# A common representation of time across visual and auditory modalities

**DOI:** 10.1101/183426

**Authors:** Louise C. Barne, João R. Sato, Raphael Y. de Camargo, Peter M. E. Claessens, Marcelo S. Caetano, André M. Cravo

**Author notes:** **Corresponding author**, André M. Cravo, Centro de Matemática Computação e Cognição, Universidade Federal do ABC, Rua Arcturus, 03. Bairro Jardim Antares. São Bernardo do Campo - SP - Brasil. CEP 09606-070.

## Abstract

Humans’ and non-human animals’ ability to process time on the scale of milliseconds and seconds is essential for adaptive behaviour. A central question of how brains keep track of time is how specific temporal information across different sensory modalities is. In the present study, we show that encoding of temporal intervals in auditory and visual modalities are qualitatively similar. Human participants were instructed to reproduce intervals in the range from 750 ms to 1500 ms marked by auditory or visual stimuli. Our behavioural results suggest that, although participants were more accurate in reproducing intervals marked by auditory stimuli, there was a strong correlation in performance between modalities. Using multivariate pattern analysis in scalp EEG, we show that activity during late periods of the intervals was similar within and between modalities. Critically, we show that a multivariate pattern classifier was able to accurately predict the elapsed interval, even when trained on an interval marked by a stimulus of a different sensory modality. Taken together, our results suggest that, while there are differences in the processing of intervals marked by auditory and visual stimuli, they also share a common neural representation.

## Introduction

The ability to estimate time is essential for humans and non-human animals to interact with their environment (Buhusi and Meck, 2005; Mauk and Buonomano, 2004; Merchant et al., 2013a). Intervals in the range of hundreds of milliseconds to seconds are critical for sensory and motor processing, learning, and cognition (Buhusi and Meck, 2005; Mauk and Buonomano, 2004; Merchant et al., 2013a). However, the mechanisms underlying temporal processing in this range are still largely discussed. A central unanswered question is whether temporal processing depends on dedicated or intrinsic circuits (Ivry and Schlerf, 2008). Dedicated models propose that temporal perception depends on central specialised mechanisms, as an internal clock, that create a unified perception of time (Ivry and Schlerf, 2008). This class of models can account for behavioural findings such as correlations in performance for some temporal tasks (Keele et al., 1985) and the observation that learning to discriminate a temporal interval in one sensory modality can sometimes be transferred to other modalities (Bueti and Buonomano, 2014).

Intrinsic models of time propose that a variety of neural circuits distributed across the brain are capable of temporal processing. One of the most known examples is the state-dependent network - SDN (Mauk and Buonomano, 2004). Within this framework, neural circuits can take advantage of the natural temporal evolution of its states to keep track of time (Mauk and Buonomano, 2004). One of the main advantages of such models is that they can explain the known differences of temporal processing across sensory modalities (van Wassenhove, 2009) and that learning a specific interval does not commonly improve temporal performance in other intervals (Bueti and Buonomano, 2014).

Given that both dedicated and intrinsic views can account for some results while not explaining others, there has been an increase in interest in hybrid models, according to which local task-dependent areas interact with a higher central timing system (Merchant et al., 2013a; Wiener et al., 2011). The main advantage of hybrid models is that they can explain why performance in some timing tasks seems to be correlated across participants, while still exhibiting modality and task-related differences (Merchant et al., 2008b,a).

In humans, studies that investigate these different models employ a variety of methods, such as behavioural, neuroimaging and neuropharmacological manipulations, on healthy participants and neurological patients (Coull et al., 2011; Ivry and Schlerf, 2008; Kononowicz et al., 2016; Merchant et al., 2013a; Wiener et al., 2011). Although the high temporal resolution of EEG should in principle be optimal to track neural processing during temporal tasks, the contribution of these methods has been controversial. One of the main difficulties is the absence of a clear electrophysiological correlate of temporal processing (for a recent review see (Kononowicz et al., 2016)). This lack of electrophysiological markers makes it hard to judge, for example, whether temporal processing in different modalities share a common representation (N’Diaye et al., 2004).

In a recent study, we have shown that multivariate pattern analysis (MVPA) can reveal spatiotemporal dynamics of brain activity related to temporal processing (Bueno et al., 2017). Multivariate approaches can take advantage of small differences in the signal across electrodes that might not be detectable using classical EEG methods. These pattern recognition methods allow the assessment of whether brain states evoked by different tasks, stimuli and sensory modalities are qualitatively similar.

In the present study, we investigated whether encoding of temporal intervals in different sensory modalities is qualitatively similar. Our behavioural results suggest that, although participants are more accurate in reproducing intervals marked by auditory stimuli, there is a strong correlation over observers in performance between modalities. Critically, we show that a multivariate pattern classifier based on EEG activity can predict the elapsed interval, even when trained on an interval marked by a different sensory modality. Taken together, our results suggest that, while there are differences in the processing of intervals marked by auditory and visual stimuli, they also share a common neural representation.

## Materials & Methods

### Participants

Twenty volunteers (age range, 18-30 years; 11 female) gave informed consent to participate in this study. All of them had normal or corrected-to-normal vision and were free from psychological or neurological diseases. The experimental protocol was approved by The Research Ethics Committee of the Federal University of ABC. All experiments were performed in accordance with the approved guidelines and regulations.

### Experimental Design

The experiment consisted of a temporal reproduction task. The stimuli were presented using the Psychtoolbox v.3.0 package (Brainard, 1997) on a 20-inch CRT monitor with a vertical refresh rate of 60 Hz, placed 50 cm in front of the participant.

Each trial started with a fixation point that participants fixated throughout the trial. After a delay (1.5 s), two flashes or tones were presented, separated by a sample interval (measured between tone or flash onsets). After a random delay (1.5 to 2.5 s), volunteers were re-exposed to the same interval. After another random delay (1.5 to 2.5 s), a ready stimulus was presented to the participant, indicating the beginning of the reproduction task 1. Volunteers had to reproduce the exposed interval, initiated by the ready stimulus (RS) and ended by a stimulus caused by a button press (R1).

In auditory blocks, the tones consisted of 1000 Hz tones (100 ms duration), while the ready and end stimuli consisted of 500 Hz tones. In visual blocks, the flashes that marked the interval consisted of yellow 0.5 degrees of visual angle discs (100 ms duration), while the ready and end stimuli consisted of magenta flashes of the same size. No direct feedback was given.

The sample intervals ranged between 750 ms and 1500 ms and were uniformly distributed. Each block (visual or auditory) consisted of 120 trials. Half of the participants performed the visual block first, while the other half performed the auditory block first.

### Behavioural analysis

Events in which intervals were reproduced as longer than double the sample interval or shorter than half the sample interval were considered errors and excluded from further analyses. The proportion of errors was low for both modalities (auditory: 0.0042 ± 0.0011; maximum proportion of rejected trials per participant= 0.0167; visual: 0.0092 ± 0.0031; maximum proportion of rejected trials per participant= 0.0583)

**Figure 1.**
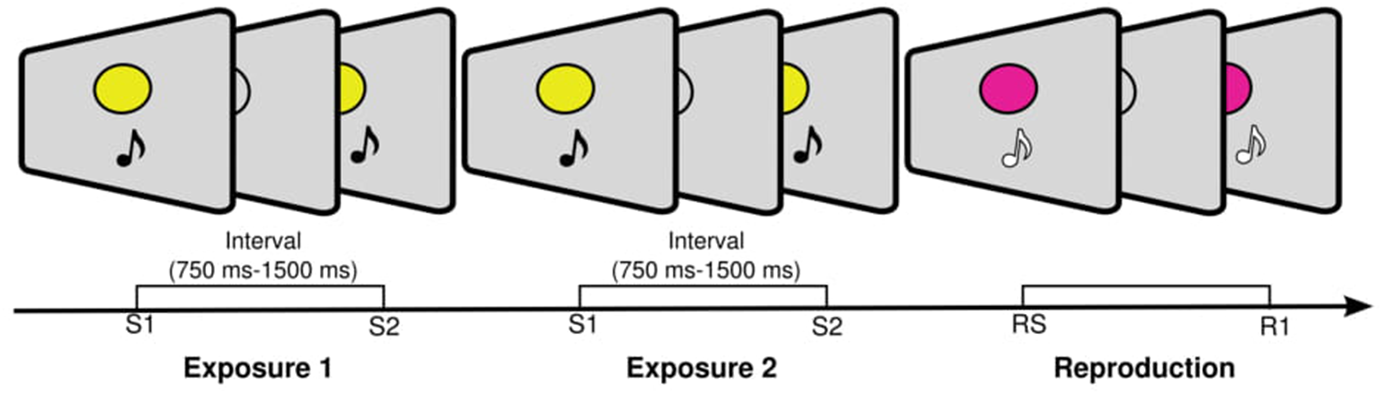
Temporal reproduction task. (A) Sequence of events during a trial. Each trial consisted of two equal empty intervals (between 750 ms and 1500 ms), marked by two stimuli. In auditory blocks, the interval was marked by two brief tones (1000 Hz, 100 ms), while in visual blocks the interval was marked by two flashes (0.5 degrees of visual angle, 100 ms). Participants were instructed to reproduce the interval at the end of each trial.

Similarly to previous studies (Cicchini et al., 2012; Jazayeri and Shadlen, 2010), the total error in the reproduction task was partitioned into two components: the average bias (BIAS) and the average variance (VAR). These two metrics are directly related to the overall mean squared error (MSE). To calculate these components, sample intervals were first binned into six equally sized bins and, for each bin, an estimate of both measures were calculated. The BIAS for each bin was calculated as the average difference between the reproduced interval and the sample intervals. The VAR for each bin was calculated as the variance of the difference between reproduced and real intervals. The final estimate of the BIAS was calculated as the root mean square of the BIAS across bins and of the VAR as the average VAR across bins (Jazayeri and Shadlen, 2010). For ease of interpretation and comparison with previous studies, the VAR values were plotted using its square root 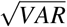.

We further calculated a regression index (RI) to index the tendency of reproduced intervals to regress towards the mean sample. This index was calculated as the difference in slope between the best linear fit on the reproduced interval and perfect performance (Cicchini et al., 2012). This measure varies from 0 (perfect performance) to 1 (complete regression to the mean, after allowing for a constant bias). The same linear fit between real and reproduced intervals was used to calculate the indifference points for both modalities. This point refers to the physical interval where durations are reproduced veridically.

To analyse the scalar property of time, we computed the slope of the generalised Weber function (García-Garibay et al., 2016; Getty, 1975; Ivry and Hazeltine, 1995). A linear regression between the variance of the reproduced intervals and the mean subjective duration squared was performed as follows:

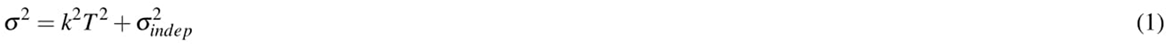

where *k* is the slope that approximates the Weber fraction and 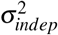 is a constant representing the time-independent component of the variability. To account for the systematic bias found in the reproduced intervals, we used an approach similar to García-Garibay et al. (2016) in which *T*^2^ was computed based on the linear fit between real and reproduced intervals.

All measures were calculated separately for each participant and condition (auditory and visual). At the group level, comparisons between the calculated parameters were done using paired t-tests (two-sided). To investigate correlations across participants for all three measures, Pearson correlations were calculated between the values of each measure in visual and auditory conditions.

### EEG recordings and pre-processing

EEG was recorded continuously from 64 ActiCap Electrodes (Brain Products) at 1000 Hz by a QuickAmp amplifier (Brain Products). All sites were referenced to FCz and grounded to AFz. The electrodes were positioned according to the International 10-10 system. Additional bipolar electrodes registered the electrooculogram (EOG).

EEG pre-processing was carried out using BrainVision Analyzer (Brain Products). All data were down-sampled to 250 Hz and re-referenced to the average activity across electrodes. For eye movement artefact rejection, an independent component analysis (ICA) was performed on filtered (Butterworth Zero Phase Filter between 0.05 Hz to 30 Hz) and segmented (−200 ms to 2000 ms relative to S1) data. For the ICA, epochs were baselined based on the average activity of the entire trial. Eye related components were identified by comparing individual ICA components with EOG channels and by visual inspection. The average proportion of rejected trials for each participant corresponded to 0.0613. For all further analyses, epochs were baseline corrected based on the period between −200 ms to 0 ms relative to S1 presentation.

### Multivariate pattern analysis - MVPA

#### Similarity between different trials and modalities

To calculate the similarity across trials we used a bootstrap approach. For each participant and comparison of interest, data from all intervals that lasted at least 1.125 seconds were divided into two groups of trials and averaged across trials per group (Bueno et al., 2017). Then, for each time point, data from each split (two-row vectors with the averaged amplitude values of all 62 electrodes) were compared using a Pearson correlation. This procedure was repeated 5000 times for each participant and Fisher transformed coefficients were averaged across permutations for each participant and comparison of interest.

At the group level, EEG-analyses were implemented non-parametrically (Maris and Oostenveld, 2007; Wolff et al., 2015) with sign-permutation tests. For each time-point, the Fisher transformed coefficients for a random half of the participants were multiplied by −1. The resulting distribution was used to calculate the p-value of the null-hypothesis that the mean value was equal to 0. Cluster-based permutation tests were then used to correct for multiple comparisons across time using 5000 permutations, with a cluster-forming threshold of p < 0.05 (Maris and Oostenveld, 2007). The sum of the values within a cluster was used as the cluster-level statistic. The significance threshold was set at p < 0.05; all tests were one-sided.

#### Decoding the elapsed interval

To investigate whether information about the elapsed interval could be decoded from activity across electrodes, we used a Naive Bayes classifier to perform a multiclass classification (Grootswagers et al., 2017). For all classifications, activity from the last 125 ms of the elapsed interval (−125 ms to 0 relative to S2) was averaged for each electrode. Data for each exposure (E1 and E2) were analysed separately. Intervals were divided into six equally sized bins (125 ms each).

#### Within modality classification

The pre-processing and decoding procedures followed standard guidelines (Grootswagers et al., 2017; Lemm et al., 2011). A leave-one-out method was used, although similar results were obtained when using a 10-fold cross-validation. For each test trial, data from all other trials were used as the training set. A principal component analysis (PCA) was used to reduce the dimensionality of the data (Grootswagers et al., 2017). The PCA transformation was computed on the training data and the components that accounted for 99% of the variance were retained. The estimated coefficients from the training data were applied to the test data. On average, the number of components used was of 37.17 ± 1.43 (mean ± s.e.m., maximum number of components=47, minimum number of components=17).

Decoding performance was summarised with Cohen’s quadratic weighted kappa coefficient (Cohen, 1968). The weighted kappa gives more weight to disagreements for categories that are further apart. It assumes values from −1 to 1, with 1 indicating perfect agreement between predicted and decoded label and 0 indicating chance (Cohen, 1968).

#### Between modalities classification

The between modality decoding (training the classifier on data from one modality and testing on the other) followed a similar structure as the within-modality decoding. Given that participants performed the first half of the experiment with one modality and the second with the other, the effects of non-stationarity are low (Lemm et al., 2011). Thus, data from all trials of one modality were used as the training set and tested against all trials of the other modality.

#### Correlation of decoding and behaviour

Similarly to our previous decoding analysis, activity from the last 125 ms of the elapsed interval (−125 ms to 0 relative to S2) was used. For each test trial, a Naive Bayes classifier was trained on trials that were shorter (at least 25 ms and up to 125 ms shorter) and longer (at least 25 ms and up to 125 ms longer) than that the physical duration of that given test trial. This classification was applied to all trials between 875 ms and 1375 ms, so that all trials had a similar number of training trials.

Based on this classification, trials were further classified into three categories: (S/S) when in both presentations (E1 and E2) the trial was classified as shorter; (S/L or L/S) when in only one of the two presentations the trial was classified as longer and (L/L) when in both presentation the trials were classified as longer.

To test whether classification might explain participants’ performance, we first fitted, for each participant, a linear function relating the physical interval and participants’ reproduced interval. The residuals from this linear fit were stored and separated as a function of the three decoding categories mentioned above. Notice that the linear fit on the behavioural results naturally takes into account participants tendency to regress towards the mean. Thus, this analysis allows isolating if trials that were encoded as shorter or longer were reproduced as shorter or longer than participants’ average reproduction for that interval duration.

### Statistical analysis

All t-tests, correlations, ANOVAs and effect sizes (Cohen’s d for t-tests and omega-squared for ANOVAs) were calculated using JASP (JASP Team, 2017). The p-values were adjusted, whenever appropriate, for violations of the sphericity assumption using the Greenhouse–Geisser correction.

## Results

### Behaviour

Following previous studies (Cicchini et al., 2012; Jazayeri and Shadlen, 2010), temporal errors were partioned into two components: the average difference between the reproduction times and the sample interval, henceforth called BIAS, and the corresponding variance of the difference between reproduced and sample intervals, abbreviated below as VAR. Our results showed that participants had similar BIAS for both modalities (*BIAS_auditory_* = 0.115 ± 0.009s, *BIAS_visual_* = 0.121 ± 0.010s, mean ±s.e.m., paired t-test, *t*_19_ = −0.634, *p* = 0.534, d=0.142). However, variance was smaller for intervals marked by auditory targets (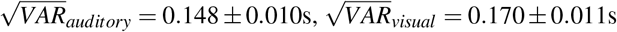, paired t-test, *t*_19_ = −2.789, *p* = 0.012, d=0.624).

As can be seen in Figure 2B, in both modalities there was a strong central tendency effect, characterised by long intervals being underestimated and short durations being overestimated (Cicchini et al., 2012; Jazayeri and Shadlen, 2010; Lejeune and Wearden, 2009; Pérez and Merchant, 2018). To measure the strength of this effect, we calculated a regression index (RI) (Cicchini et al., 2012). This index is calculated as the difference in slope between the best linear fit on the reproduced interval and perfect performance. It varies from 0 (perfect performance) to 1 (complete regression to the mean, after allowing for a constant bias). The estimated RIs suggest that participants had a stronger central tendency effect in visual blocks, as shown by higher RIs (*RI_auditory_* = 0.346± 0.036, *RI_visual_* = 0.430 ± 0.032, paired t-test, *t*_19_ = −3.152, *p* = 0.005, d=0.705).

**Figure 2.**
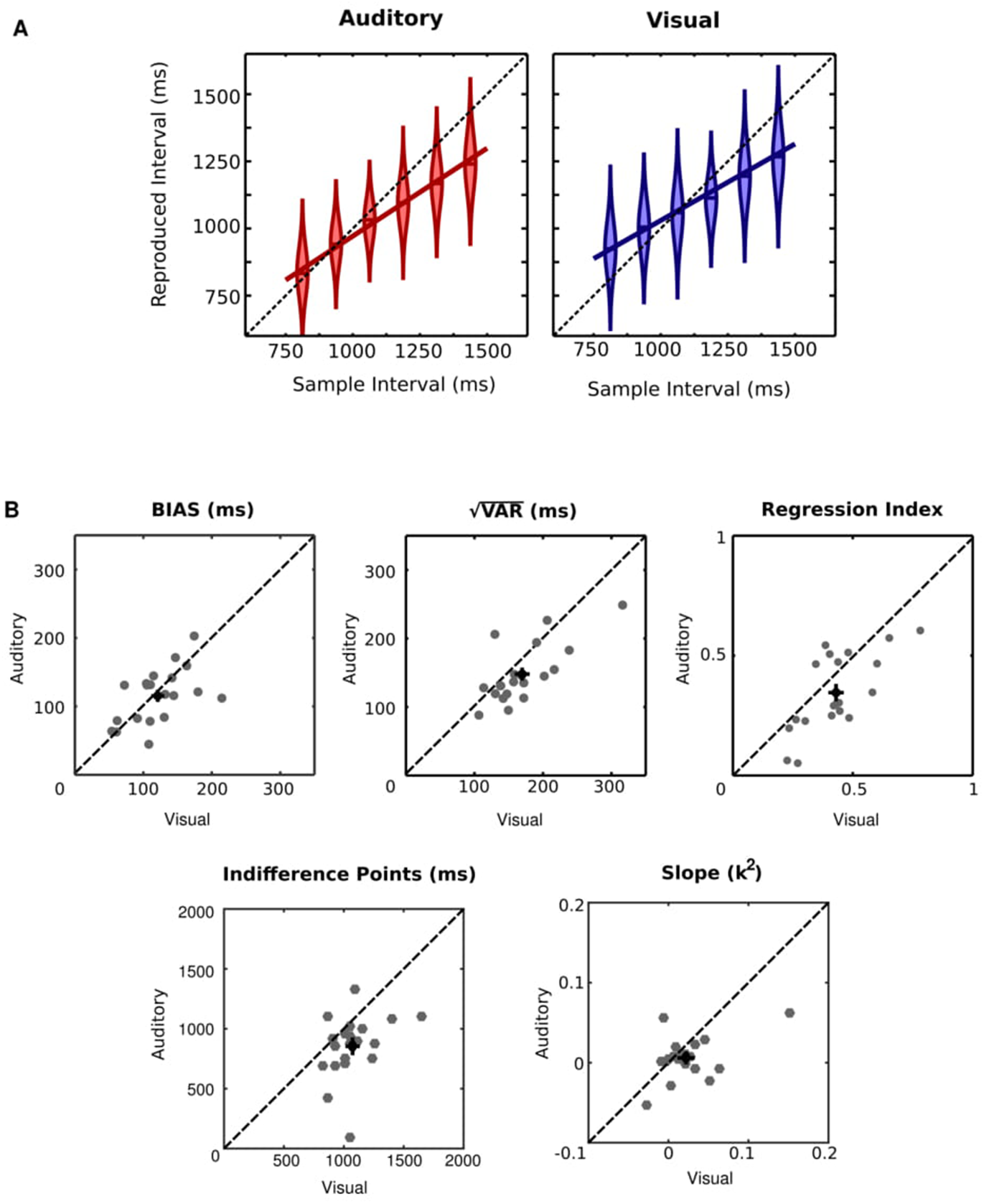
Behavioural performance (A) Average reproduction times across different intervals and conditions. Sample intervals were binned into six equally sized (125 ms) categories. Violin plots show the probability density across participants for each bin. Markers represent the median. Participants showed a central tendency effect with shorter intervals being overestimated and longer intervals being underestimated. (B) Summary statistics for performance. The BIAS (upper left panel) represents the average difference between the reproduction times and the sample interval. The VAR (upper central panel) represents the variance of the reproduced intervals. The Regression Index (upper right panel) shows the strength of the tendency to regress to the mean. The indifference points (lower left panel) represents the estimated physical interval in which participants reproduce the intervals veridically. The slope of the generalised Weber function (lower right panel) approximates the Weber fraction. Participants were, in general, better in reproducing auditory than visual intervals. There was also a correlation between visual and auditory modalities across participants for the majority of measures.

We further calculated the indifference points for both modalities. As can be seen in Figure 2B, indifference points were shorter for auditory intervals than for visual intervals (*IndifferencePoint_auditory_* = 0.854 ± 0.059, *IndifferencePoint_visual_* = 1.07 ± 0.044, paired t-test, *t*_19_ = −3.648, *p* = 0.002, d=0.816).

We used a slope analysis to analyse the scalar property of time (García-Garibay et al., 2016; Getty, 1975; Ivry and Hazeltine, 1995). A linear regression between the variance of the reproduced intervals and the mean subjective duration was used to calculate the generalised Weber function (García-Garibay et al., 2016; Getty, 1975; Ivry and Hazeltine, 1995). We found a lower slope for auditory than for visual intervals (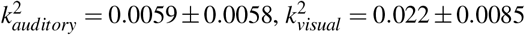, paired t-test, *t*_19_ = −2.082, *p* = 0.051, d=0.465)

Although participants were, in general, more accurate for auditorily marked compared to visually marked intervals, there was also a strong correlation in how well participants performed for both modalities, in agreement with previous findings (Merchant et al., 2008b; Stauffer et al., 2012). As shown in Figure 2B, there was a strong correlation between performance in auditory and visual blocks across participants as measured in their BIAS (Pearson product-moment correlation: *r*_20_ = 0.545, *p* = 0.013), 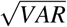 (*r*_20_ = 0.722, *p* < 0.001), Regression Index (*r*_20_ = 0.701, *p* < 0.001) and slopes (*r*_20_ = 0.475, *p* = 0.034). Indifference points were not significantly correlated across participants (*r*_20_ = 0.339, *p* = 0.144).

### EEG recordings

#### Similarity of the evoked activity for intervals of similar modalities

The evoked ERPs resembled classical results from temporal processing studies, with a strong contigent negative variation (CNV) in frontocentral electrodes for both modalities (auditory/visual) and exposures (E1/E2, Figure 3). If a given pattern of activity carries information about time, its sequence of activation must be consistent across different trials. Thus, in a first step, we investigated whether the temporal pattern of EEG activity was similar in different trials within the same modality (auditory or visual). To have a sufficient number of trials, EEG activity was epoched relative to S1 for all trials that had an interval equal or longer than 1125 ms. These trials were then randomly divided into two groups of trials and averaged per group. The similarity of activity across sensors for the different splits was quantified by means of a Pearson correlation coefficient. This procedure was repeated 5000 times for each participant and Fisher transformed coefficients were averaged across permutations for each participant.

**Figure 3.**
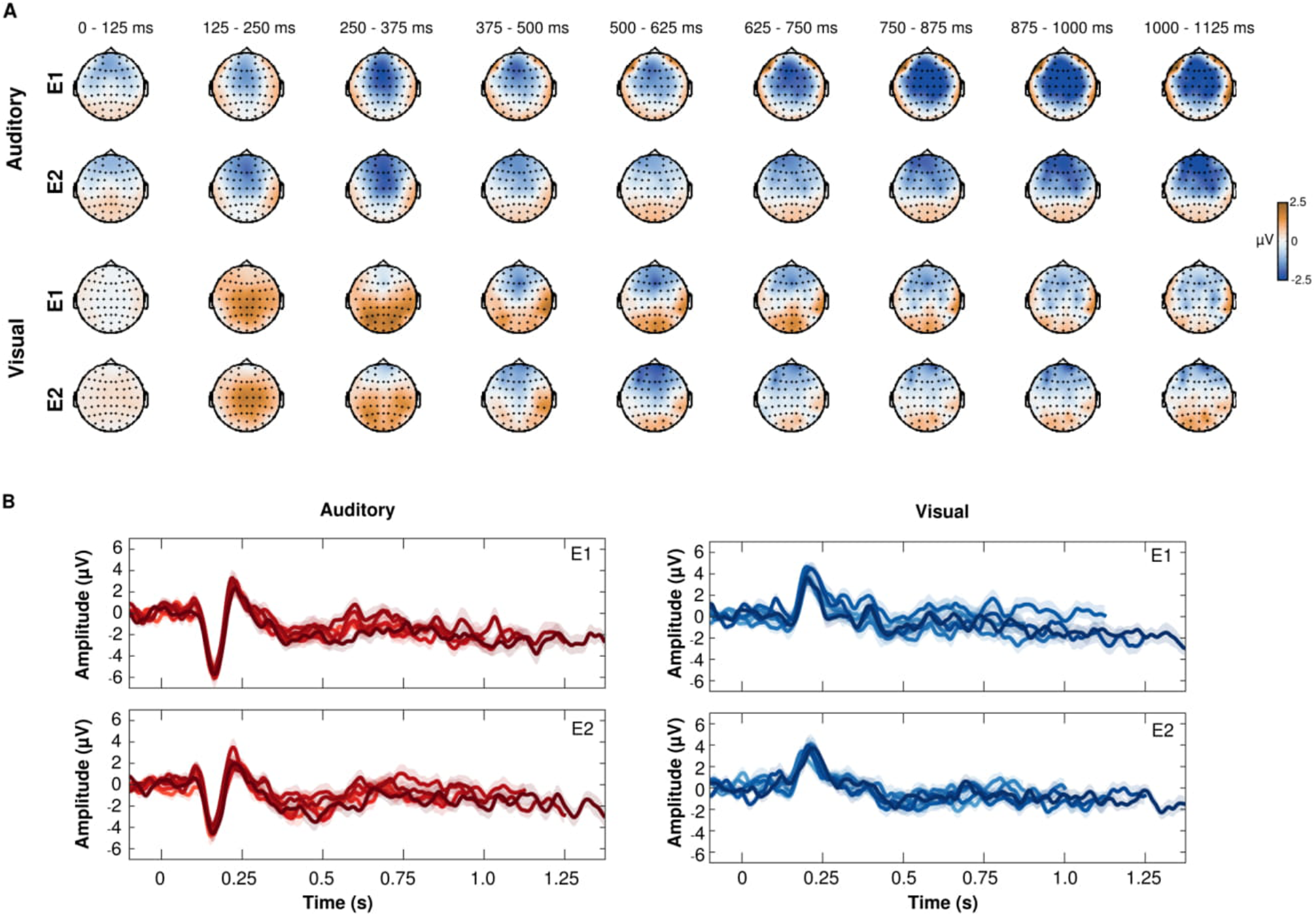
(A) Topographies for intervals longer than 1.125 seconds for both modalities and exposures. Each topography shows the average activity in 125 ms temporal bins, from 0 s to 1.125 s relative to S1 (B) Event-related potentials (mean ±s.e.m.) in fronto-central electrodes (FC2,FCz,FC1,C1,Cz,C2) for each of the six binned tested durations (bin edges:750ms-875ms; 875ms-1000ms; 1000ms-1125ms; 1125ms-1250ms; 1250ms-1375ms and 1375ms-1500ms). Both panels show epochs segmented relative to S1, in both exposures (E1 and E2). Darker (lighter) colours represent longer (shorter) intervals.

As shown in Figure 4A, for trials of the same modality, there was strong similarity across trials that started shortly after the first stimuli that marked the interval (S1). In auditory blocks, a statistically significant temporal cluster emerged 85 ms after S1 for the first exposure (E1, cluster-p < 0.001) and after 120 ms for the second (E2, cluster p < 0.001). In both conditions, the cluster remained throughout the interval (Figure 4A, left panel). In visual blocks, a statistically significant cluster emerged 96 ms after S1 for E1 (cluster p < 0.001) and after 84 ms for E2 (cluster p < 0.001). Similarly to auditory blocks, the clusters also remained significant throughout the interval (Figure 4A, middle panel).

**Figure 4.**
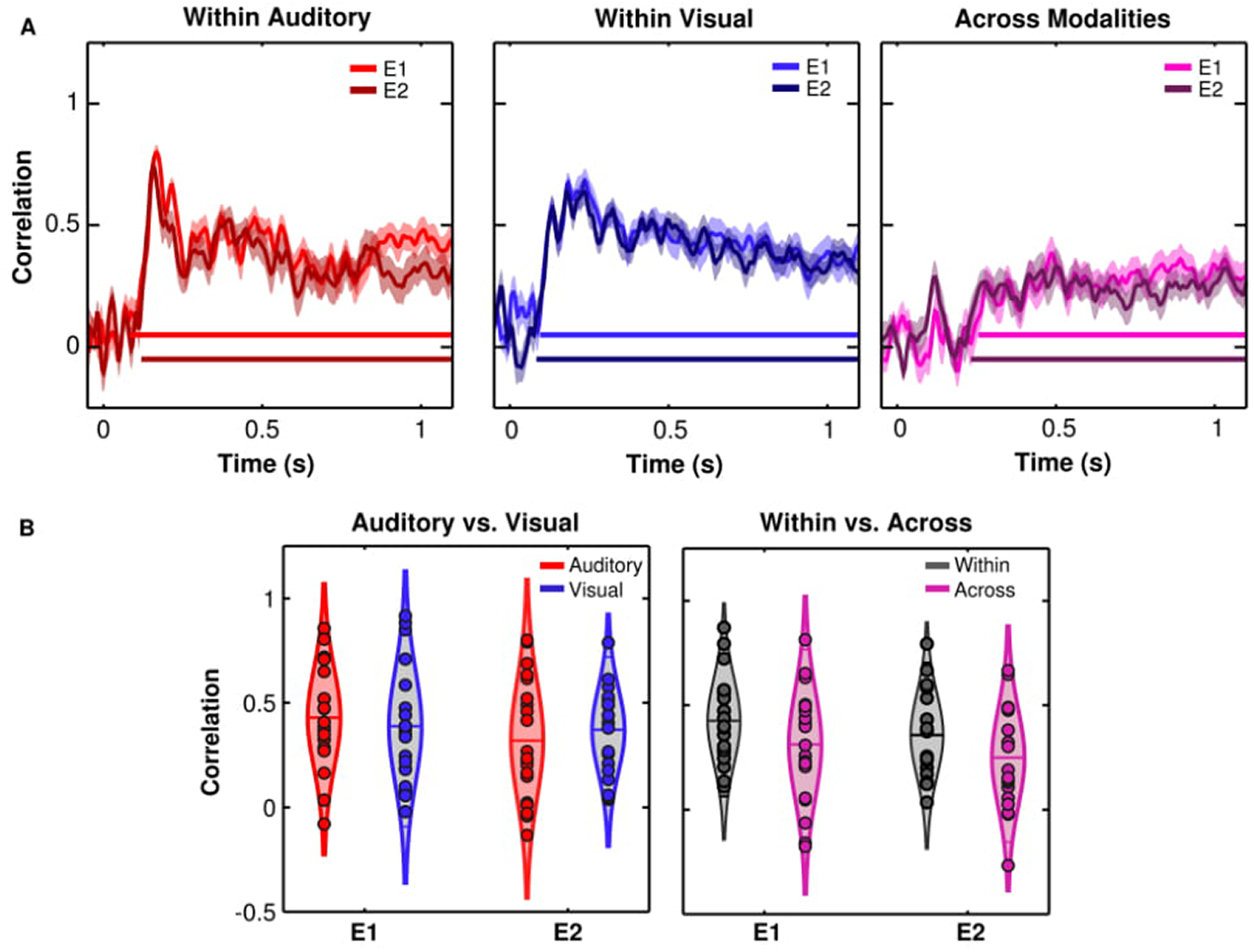
Pattern similarity across trials of same or different modalities. (A) The similarity index (mean ±s.e.m. of estimated Pearson correlation) shows the consistency of the pattern of activity across trials within auditory blocks (left panel), visual blocks (middle panel) and across modalities (right panel). In all panels, lighter/darker lines represent similarity for the first (E1) and second exposure (E2) in each trial. Continuous lines at the bottom of each panel indicate periods of significant similarity across trials. (B) Comparison of pattern similarity for trials of the same (left panel) or different (right panel) modalities. Violin plots show the probability density across participants. Line markers represent the median and interquartile range. Circles inside the violin plots show data from all participants.

In a next step, we investigated whether similarity was modulated by sensory modality (auditory/visual) or exposure within the trial (E1/E2). Similarity indexes over the final 400-ms period of the selected trials (725 ms to 1125 ms) were averaged for each modality (visual/auditory) and exposure (E1/E2) and submitted to a repeated measures ANOVA (Figure 4B, left panel). There was no significant modulation by modality (F1,19 = 0.004, p = 0.948, ***ω***^2^ = 0), exposure (*F*_1,19_ = 1.973, *p* = 0.176, ***ω***^2^ = 0.044) or interaction (*F*_1,19_ = 0.118, *p* = 0.735, ***ω***^2^ = 0).

#### Similarity of the evoked activity for intervals across modalities

As expected, the pattern of activity during the interval was similar for trials of the same sensory modality. However, if there is a common representation of time across different modalities, the pattern of activity for intervals of different modalities should also be consistent. To test this possibility, a similar analysis as described above was performed with EEG activity from intervals marked by different modalities. Similarity across trials emerged later for these comparisons. Significant clusters emerged at 256 ms after for E1 trials (cluster p < 0.001) and at 232 ms after S1 for E2 trials (cluster p < 0.001, Figure 2A, right panel). Thus, although the pattern of EEG activation was similar across visual and auditory intervals, this similarity started at later moments than for within-modality comparisons.

In a next step, the magnitude of the similarity when comparing trials of same and different modalities was tested. Similarity indexes over the last 400 ms of the investigated intervals (725 ms to 1125 ms) were averaged for each exposure (E1/E2) and modality (within/across) and submitted to a repeated measures ANOVA (Figure 2B, right panel). Similarity for intervals of the same modality was significantly stronger than for different modalities (*F*_1,19_ = 28.443, *p* < 0.001, ***ω***^2^ = .567). There was no significant effect of exposure (*F*_1,19_ = 1.824, *p* = 0.193, ***ω***^2^ = 0.038) or of the interaction of factors (*F*_1,19_ = 0.057, *p* = 0.813, ***ω***^2^ = 0).

#### Decoding time from EEG signals

Given the strong consistency of activity for intervals within and across modalities, we further investigated whether this pattern might carry information about the elapsed time. In a first analysis, we focused on intervals of the same sensory modality.

Activity from the final period of each interval (−125ms to 0 relative to S2) was binned into six equally spaced periods (bin edges: 750ms-875ms; 875ms-1000ms; 1000ms-1125ms; 1125ms-1250ms; 1250ms-1375ms and 1375ms-1500ms). A Naive Bayes classifier was used to decode from which of the six binned intervals activity from each trial came from (Gouvêa et al., 2015; Grootswagers et al., 2017). A weighted Kappa coefficient was used to quantify the agreement between decoded and real interval. This coefficient ranges from 1 (perfect agreement) to −1 (complete disagreement), where 0 indicates chance levels (Cohen, 1968).

As shown in Figure 5, there was a strong and statistically significant agreement between predicted and real intervals in both auditory (E1: *Kappa_auditory_* = 0.217 ± 0.026, t-test against zero, *t*_19_ = 8.480, p< 0.001, d=1.896; E2: *Kappa_auditory_* = 0.237 ± 0.024, t-test against zero, *t*_19_ = 9.887, p< 0.001, d=2.211) and visual blocks (E1: *Kappa_visual_* = 0.225 ± 0.021, t-test against zero, *t*_19_ = 10.623, p< 0.001, d=2.375; E2: *Kappa_visual_* = 0.224±0.026, t-test against zero, *t*_19_ = 8.598, p< 0.001, d=1.923).

**Figure 5.**
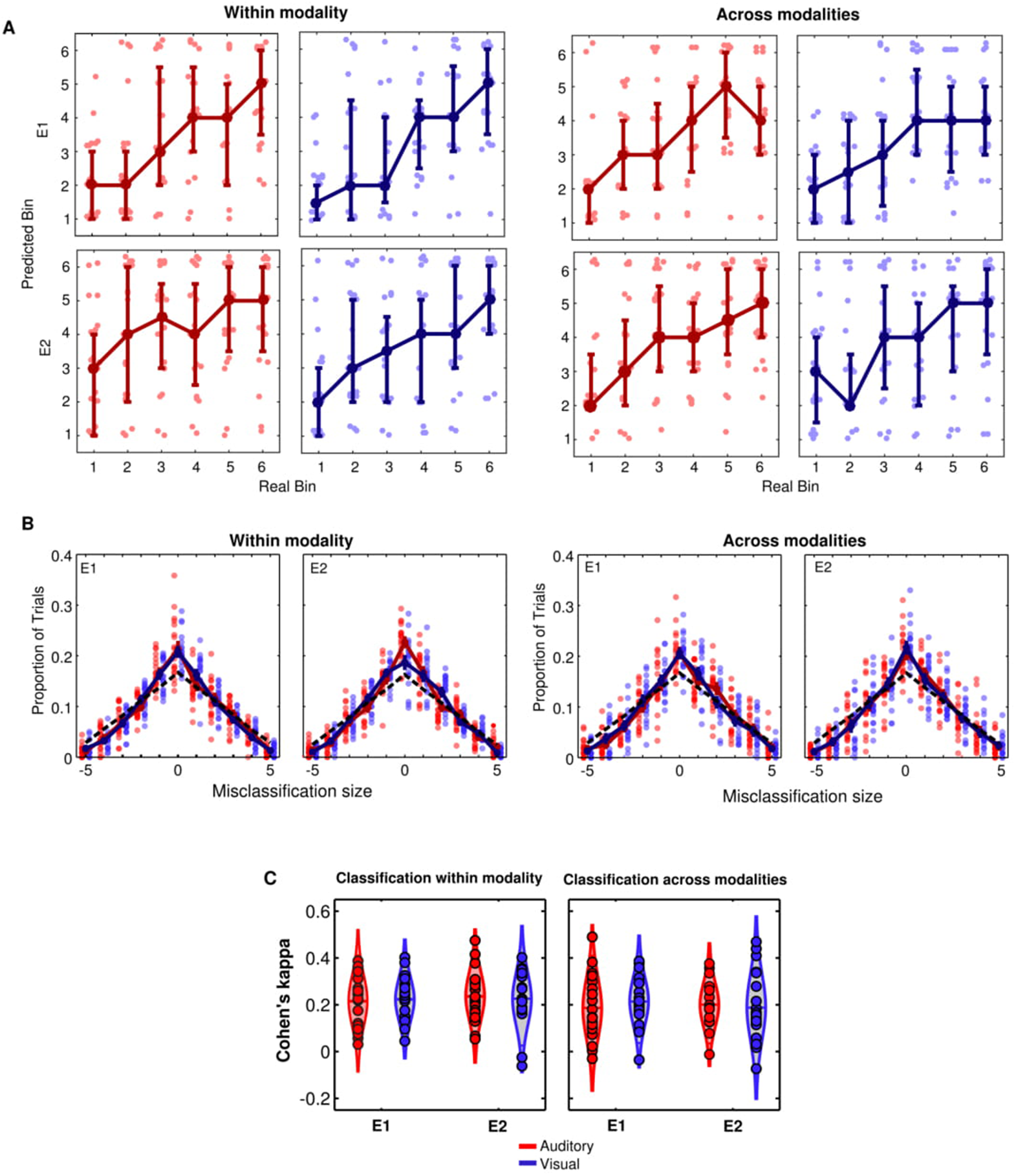
MVPA performance for interval classification within and between sensory modalities. (A) Observed versus predicted intervals by the Naive Bayes classifier across exposures and modalities. Each circle shows, for each participant and tested bin, the most common output of the classifier (mode). The continuous line shows the median across participants and error bars show the lower quartile (25th percentile) and upper quartile (75th percentile) across participants. (B) Classification performance of the Naive Bayes classifier when trained within the same sensory modality (upper panels) or between sensory modalities (lower panels). Each plot shows the proportion of trials in which the classifier was able to decode the interval correctly (when the error in the x-axis equals zero) and the proportion of trials in which different levels of errors were observed. The black dashed line represents the expected pattern of errors if the classifier were at chance levels. Left/right panels show accuracy for the first/second exposure (E1/E2) in each trial. Error bars and continuous lines show mean error (mean ±s.e.m.) across participants. Each circle shows data from one participant. (C) Agreement between decoded and real interval indexed by Cohen’s kappa. The left panel shows classification performance when test trials were of the same modality as the training trials while right panels show performance when the test trials were of a different modality than the training trials. Violin plots show the probability density across participants; line markers represent the median. Circles inside the violin plots show data from each participant.

Given that the elapsed interval could be decoded from EEG activity and that this activity was similar across different sensory modalities, we investigated whether similar decoding performance could be achieved across sensory modalities. A similar Naive Bayes classifier was used as described above. However, to test whether decoding could be generalised across sensory modalities, each test trial was classified based on the activity of intervals of the other modality.

Once again, there was strong agreement between predicted and real intervals, as measured by weighted Kappa coefficients. This was true when auditory intervals were decoded using visual intervals as training sets (E1: *Kappa_auditory_* = 0.185 ± 0.030, t-test against zero, *t*_19_ = 6.137, p< 0.001, d=1.372; E2: *Kappa_auditory_* = 0.199 ± 0.022, t-test against zero, *t*_19_ = 9.0627, p< 0.001, d=2.026) as well as for when visual intervals were decoded using auditory intervals as training sets (E1: *Kappa_visual_* = 0.216 ± 0.024, t-test against zero, *t*_19_ = 9.020, p< 0.001, d=2.017; E2: *Kappa_visual_* = 0.190 ± 0.033, t-test against zero, *t*_19_ = 5.800, p< 0.001, d=1.297).

A repeated measures ANOVA was used on the estimated Kappas to test whether decoding performance depended on the modality to be predicted (auditory or visual), on the exposure (E1 or E2) and on whether it was trained on the same or different modality (within modality/across modalities). There was a trend of decoding performance being higher when trained within rather than between modalities (*F*_1,19_ = 3.502, *p* = .077, ***ω***^2^ = 0.106). There were no significant effects of modality to be predicted (*F*_1,19_ = 0.131, *p* = .721, ***ω***^2^ = 0) or of exposure (*F*_1,19_ = 0.013, *p* = .909, ***ω***^2^ = 0). None of the interactions were significant (modality to be predicted*exposure: *F*_1,19_ = 1.814, *p* = .194, ***ω***^2^ = 0.037; modality to be predicted*trained modality: *F*_1,19_ = 0.132, *p* = .721, ***ω***^2^ = 0; trained modality*exposure:*F*_1,19_ = 0.224, *p* = .642, ***ω***^2^ = 0; trained modality*exposure*modality to be predicted:*F*_1,19_ = 0.114, *p* = .739, ***ω***^2^ = 0).

#### Decoded interval and behavioural performance

We further investigated whether the pattern of EEG activity during the encoding of intervals was associated with the reproduced interval. Specifically, we tested if trials in which activity during encoding was more similar to intervals that were shorter/longer than its real duration were followed by systematic under/overestimation of the reproduced interval.

Similarly to our previous analysis, the activity of the end of each interval (−125ms to 0 relative to S2) was used. For each test trial, the Naive Bayes classifier was trained exclusively in intervals that were physically longer or shorter than the interval presented in that trial. The output of the classifier for each trial can be thought of as a forced judgement of whether, in a given trial, the EEG activity is more similar to intervals longer or shorter than that interval. Because each interval was presented twice during encoding (E1 and E2), this procedure was performed for each presentation separately. Based on the classifier outputs, trials were classified into three categories: (S/S) when in both exposures (E1 and E2) the trial was classified as shorter, (S/L or L/S) when in one of the two exposures the trial was classified as longer, and (L/L) when in both exposures the trials were classified as longer.

To test whether these categories were correlated with performance, we focused on the residuals of the linear regression between presented and reproduced interval. These residuals reflect fluctuations in the reproduced interval that were above or below participants’ mean reproduced interval for that given duration. Residual behaviour for each participant was separated according to the three categories and submitted to a repeated measures ANOVA with category (S/S; S/L or L/S; L/L), modality (auditory or visual) and trained modality (within or across) as factors. There was a main effect of category (*F*_1,19_ = 7.231, *p* = .003, ***ω***^2^ = .233). A post-hoc analysis showed a linear trend between predicted category and residual behaviour (*F*_1,19_ = 10.898, *p* = .004), suggesting an association between whether the interval was classified as shorter or longer and the reproduced interval (Figure 6). The other main effects were not significant (modality: *F*_1,19_ = 0.018, *p* = 0.893, ***ω***^2^ = 0; trained modality: *F*_1,19_ = 0.204, *p* = 0.657, ***ω***^2^ = 0), nor the interactions (category*modality: *F*_1,19_ = 0.373, *p* = 0.691, ***ω***^2^ = 0; category*trained modality:*F*_1,19_ = 0.183, *p* = 0.777, ***ω***^2^ = 0; trained modality*modality: *F*_1,19_ = 0.066, *p* = 0.799, ***ω***^2^ = 0; category*trained modality*modality: *F*_1,19_ = 0.104, *p* = 0.857, ***ω***^2^ = 0).

**Figure 6.**
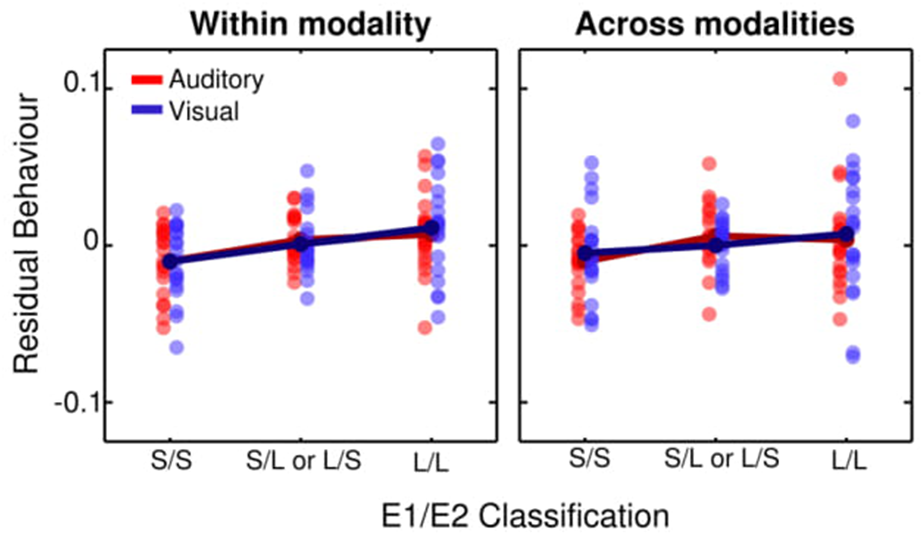
Correlation between MVPA classification and performance. Both panels show the residual reproduced interval as a function of test trials being classified as short in both E1 and E2 (S/S); as short in one of the exposures and as long in the other (S/L or L/S) or as long in both E1 and E2 (L/L). Each circle shows data from one participant and continuous lines show mean residual across participants. The left panel shows results when the test data were classified based on trials of the same modality. The right panel shows results from when the test data were classified based on trials of a different modality.

## Discussion

In this study, we investigated whether there is a common representation of time for auditory and visual modalities. The behavioural results showed that, although performance was better for auditory intervals than for visual intervals, there was a strong correlation in performance across participants. The electrophysiological findings showed that activity elicited by intervals was consistent shortly after its onset when of the same modality, and later when of different modalities. Finally, a classifier based on multivariate activity across electrodes was able to decode temporal intervals within and between modalities and was correlated with behavioural performance.

Our behavioural results are in agreement with the general finding that temporal acuity is better for auditory than for visual intervals (Merchant et al., 2008b,a; van Wassenhove, 2009). Differences in performance between sensory modalities have usually been considered as evidence for distributed temporal processing. On the other hand, and also in agreement with previous behavioural findings, we found a strong correlation in performance between modalities (Merchant et al., 2008b,a; Stauffer et al., 2012). In our specific experiment, one could argue that the behavioural correlations between performances in different modalities are being driven by the motor task and not by a common encoding of time across modalities. However, given that performance differed between auditory and visual intervals, it seems unlikely that the observed correlations are due exclusively to the motor output of the task. Moreover, previous behavioural studies that found correlations across different modalities and tasks have used a wide variety of tasks, including purely perceptual judgements (Merchant et al., 2008b,a; Stauffer et al., 2012). Thus, our results suggest that the encoding of these intervals might being taking place in a common representation across visual and auditory modalities. Nevertheless, it would be interesting for future studies to address whether this common representation of time is also task-dependent.

Associations in performance across modalities and temporal tasks have been used as evidence for the hypothesis that different modalities share a central mechanism for temporal processing (Merchant et al., 2013a). An influential view is that, while temporal processing of short intervals (on the scale of a few hundred milliseconds) can be mediated by perceptual mechanisms, longer intervals (on the scale of seconds and minutes) might depend on later, cognition-based mechanisms (Lewis and Miall, 2003; Rammsayer et al., 2015; Stauffer et al., 2012). In the specific case of different sensory modalities, it has been suggested, based on behavioural findings, that temporal processing goes through a gradual transition of a modality-specific to a cognitive amodal timing mechanism(Rammsayer et al., 2015; Stauffer et al., 2012). We provide electrophysiological evidence that is consistent with this view. In our findings, the pattern of electrophysiological activity was consistent for the early moments when comparing intervals of the same modality. For intervals of different modalities, there was also a consistent pattern of activity that emerged later in the interval. While the existence of a hard boundary between a local, modality-specific, and a shared representation of time is unlikely (Buonomano et al., 2009), our results are in agreement with recent findings that suggest that this transition might take place in the range of hundreds of milliseconds (Buonomano et al., 2009; Karmarkar and Buonomano, 2007; Merchant et al., 2013a) and not in the range of seconds as has previously been suggested. Although we found a common representation of time in later intervals, this finding does not necessarily suggest that temporal processing is not going on in modal regions. Rather, modality-specific timing information could be processed locally and passed to common shared regions.

The transition of a local sensory-dependent pattern of activity to a commonly shared representation of time is consistent with hybrid models of temporal processing, in which there is a transition between sensory encoding of timing information and a later representation of time in the motor system to control behaviour (Merchant et al., 2013a; Merchant and Yarrow, 2016; Wiener et al., 2011). Based on neuroimaging and invasive electrophysiological recordings in non-human animals, it has been proposed that this higher core-timing network depends on structures of the cortico-thalamic-basal ganglia circuit (Buhusi and Meck, 2005; Coull et al., 2011; Merchant et al., 2013a), such as the supplementary motor area (SMA) (Merchant et al., 2013a, 2015). Recent studies have suggested that in several of these regions time is encoded by means of a population code (Bakhurin et al., 2016; Crowe et al., 2014; Gouvêa et al., 2015; Merchant et al., 2013b; Matell et al., 2011). As previously mentioned, how this population code is translated to different EEG signals is a topic of discussion. However, here we show that multivariate decoding analysis might overcome these limitations and reveal important properties of temporal processing. In our results, a multivariate classifier was able to decode the elapsed interval when it was marked by the same or by a different modality as the training data. This results suggests that, at least for this late period, the representation of time is qualitatively similar in different sensory modalities.

In a previous study, we have already shown that using multivariate pattern analysis on the EEG signal can reveal activity related to temporal processing (Bueno et al., 2017). However, in that experiment, MVPA was applied while participants performed a temporal categorisation task in which they had to judge whether a certain duration was shorter, equal or longer than a reference. Thus, it was possible that the pattern analysis was picking up activity related to decisional processing or to particular strategies adopted by the participants (Bueno et al., 2017). Similarly, a study that also investigated whether the pattern of EEG activity during temporal judgements was consistent across vision and audition focused its analysis during the interval to be judged (N’Diaye et al., 2004). Here, on the other hand, the classifier was trained during the encoding of an interval that had to be reproduced later, reducing the possibility of a motor or decisional contamination of the signal. Nevertheless, the pattern of activity during the encoding was correlated with the reproduced interval, suggesting, once again, that this activity is related to temporal processing.

Taken together, our results suggest that, while there are differences in the processing of intervals marked by auditory and visual stimuli, they also share a common late neural representation. In addition, they show how the analysis of multivariate patterns can bring about important advances in our understanding of temporal processing.

## Acknowledgements

This work was supported by the Fundação de Amparo à Pesquisa do Estado de São Paulo (FAPESP) research grants 13/24889-7 and 14/08389-7 and by Conselho Nacional de Pesquisa (CNPq) research grant 122942/2014-0. The authors would like to thank Nicholas E. Myers and the members of the Timing and Cognition Laboratory at UFABC (http://neuro.ufabc.edu.br/timing/) for useful discussions and suggestions on earlier versions of this manuscript.

**Author contributions statement**: L.C.B., M.S.C, P.M.E.C and A.M.C. conceived the experiment. L.C.B. performed the experiments. All authors analysed the data, wrote and reviewed the manuscript.

